# Chromatin-modifying enzymes as modulators of nuclear size during lineage differentiation

**DOI:** 10.1101/2022.05.15.491991

**Authors:** Lingjun Meng

## Abstract

The mechanism of nuclear size determination and alteration during normal lineage development and cancer pathologies which is not fully understood. As recently reported, chromatin modification can change nuclear morphology. Therefore, we screened a range of pharmacological chemical compounds that impact the activity of chromatin-modifying enzymes, in order to get a clue of the specific types of chromatin-modifying enzymes that remarkably effect nuclear size and shape. We found that interrupted activity of chromatin-modifying enzymes is associated with nuclear shape abnormalities. Furthermore, the activity of chromatin-modifying enzymes perturbs cell fate determination in cellular maintenance and lineage commitment. Our results indicated that chromatin-modifying enzyme regulates cell fate decision during lineage differentiation and is associate with nuclear size alteration.

**Significance:** Here we described for the first time the modulation of chromatin-modifying enzymes remarkably effects nuclear shape, and perturbs cell fate determination in cellular maintenance and lineage fate commitment in normal stem cells and leukemia cancer cells. We found that the irregularities of nuclear contour were highly related to pharmacological inhibition of chromatin-modifying enzyme activity. After manipulating a histone demethylase named GASC1, we found that upregulation of GASC1 impairs the differentiation of hESCs to terminally differentiated neural cells. Moreover, upregulation of GASC1 impairs the proliferation of leukemia cells due to cell cycle arrest in the G0/G1 phase and indicates misshapen nuclei. This study suggested that chromatin-modifying enzyme regulates the nuclear contour related cell fate decision also including cancer cell fate determination.

## Introduction

As reported, nuclear volume is an important morphological parameter that can change during cancer progression (1). After confocal microscopy was discovered and broadly used to observe the detailed structure of specific objects within the cell, the most fascinated cellular organelle - nucleus was researched more carefully. Recent studies have also highlighted that nuclear shape can influence fate decisions (2). Embryonic stem cells (ESCs) differentiation is accompanied by an increase in chromatin condensation (3) leading to an increase in nuclear stiffness (4) and a decrease in nuclear size. As known, the nuclear size of stem cells is extremely bigger than terminal differentiated cells. Cell differentiation is corresponding with chromatin marks reset, which raises the question of whether chromatin’s contribution to nuclear mechanics affects nuclear morphology as well as cell fate. It is therefore critical to identify the contributions of various chromatin-modifying enzymes that are relevant to cell fate decisions, and consequently display nuclear size alteration.

Nuclear architecture is impacted via chromosome structure which is determined by histone modifications. The cellular differentiation from stem cells to differentiated cells which is corresponding with nuclear size reduction as well as chromosome condense. Histone demethylases have also been reported to be critical enzymes that regulate the balance between self-renewal and differentiation in hESCs (human embryonic stem cells) (5–8). Accordingly, we broadly utilized chromatin-modifying enzymes inhibitors to treat stem cells such as hESCs, NPCs (neural progenitor cells), and later cancer cells such as leukemia cells. We demonstrated that the irregularities of nuclear contour were highly related to pharmacological inhibition of chromatin-modifying enzyme activity. As known, upregulation or downregulation of histone demethylases is observed in various types of cancer and is associated with higher rates of malignancy and relapse after treatment (9). In particular, GASC1 (gene amplified in squamous cell carcinoma, also known as KDM4C, a histone demethylase) promotes chromosomal instability and transcription initiation,therefore, plays a causative role in cancer cells driven by its dysregulation (10). GASC1 was found aberrant expression in esophageal squamous carcinomas (11), medulloblastomas (12), glioma (13), breast cancer (14), and prostate cancer (15). In addition, GASC1 plays an essential role in the transition of mouse ESCs (mESCs) to endothelial cells (16). Consistently, we demonstrated that upregulation of GASC1 impairs the differentiation of undifferentiated hESCs to terminally differentiated neural cells. Moreover, leukemia cells with upregulated GASC1 expression lost their proliferation ability, show cell cycle arrest in the G0/G1 phase, and appear misshapen nucleus. Taken together, epigenetic perturbations are likely to be the cause for misregulation of nuclear size and of certain cancer cell fates (17).

## Result

### The nuclear size is associated with cell fate decision and chromatin-modifying enzyme activity

Size and shape are distinctive aspects of nuclear structure. Nuclear size impacts cell function, and is altered according to the specific status of cell types during cell development (18). It is unclear which modulators effect the altered nuclear size during cell fate transformation. To investigate the distinct nuclear size of various cell types, we employed highly efficient differentiation methods to interrogate the size change of nucleus during lineage differentiation. To differentiate NPCs from hESCs, we used two molecular compounds, SB431542 and LDN193189, to inhibit the TGF-β (transforming growth factor beta) and BMP (bone morphogenetic protein) pathways, which are also called Dual SMAD inhibitors (19). In addition, as we previously used, overexpression of Ngn2 (neural gene neurogenin-2) could converts hESCs into iN cells (induced neuronal cells) with nearly 100% yield in less than two weeks. The hESCs-derived iN cells grow long axons that conduct electrical impulses away from the neuron’s cell body (20). We utilized DAPI to stain hESCs for 5 min, then examined the dynamic change of the nuclear size via measuring the area of nucleus under a fluorescence microscope (Fig. 1*A*). Notably, it was shown that there is a reduced size of approximately 40% from hESCs to NPCs, and 24% decrease from NPCs to iN cells (Fig. 1*B*). Consistently as reported, the nuclear size of hESCs is bigger than NPCs, and the nuclear size of NPCs is bigger than iN cells. In addition, it was shown that the nuclear size of hESCs and NPCs are in a wider range than iN cells, which indicates that there is a potential of stem cells in nuclear dynamics. The different sizes of nuclei may correspond with hESCs, NPCs, and neural cells characterized with distinct unique chromatin architecture. During the reprogramming of cell identity and cell fate determination, heterochromatin plays a critical role. H3K9me3 (Tri-methylation of histone H3 lysine 9) is a histone mark associated with heterochromatin, and has emerged as a key player in repressing lineage-inappropriate genes, and shielding them from activation by transcription factors. We investigated the H3K9me3 expression in hESCs and hESC-derived NPCs by immunofluorescence staining. It was shown that there are high levels of expression of H3K9me3 in the nucleus (Fig. 1*C*). To directly address the role of chromatin in nuclear morphology, we employed a range of commercially available chemical compound inhibitors of the enzymes that modulate histone modification state to manipulate chromatin alterations to treat hESCs (Fig. 1*D* and *E*). These available inhibitors of chromatin-modifying enzymes that were represented the various functions of chromatin regulation, which helped us to investigate broadly chromatin-modifying enzymes associated with nuclear alteration. We purchased these inhibitors from tocris as the method shown. As reported, GSK-J1 is a highly potent inhibitor of the H3K27 histone demethylases JMJD3 (KDM6B) and UTX (KDM6A), likewise inhibits KDM5B, KDM5C, and KDM5A to increase H3K27me3. Histone demethylases inhibitor increased heterochromatin levels and chromatin-based nuclear stiffness. Heterochromatin is highly condensed DNA that leads to gene silencing (21). R2HG is oncometabolite produced by mutant isocitrate dehydrogenase 1 (IDH1) and IDH2, inhibits KDM4A in murine embryonic fibroblasts as well as increases H3K9me3, H3K36me3. 5-Aza is a DNA methyltransferase inhibitor that induces demethylation. GSK-LSD1 is a potent and selective lysine specific demethylase 1 (LSD1) inhibitor, increasing H3K4me3, H3K9me3, and H3K27me3. IOX1 is a histone demethylase JMJD inhibitor, increasing H3K36me3, H3K9me3, and H3K27me3. JIB04 is a pan Jumonji histone demethylase inhibitor that inhibits H3K4 demethylases, decreases H3K36me3, H3K9me3. NSC636819 is a KDM4A/KDM4B inhibitor, increasing H3K9me3. PBIT is a JARID1 (Jumonji AT-Rich Interactive Domain 1) inhibitor that increases H3K4me3. C646 is a inhibit histone acetyltransferase, which corresponds predominantly to decompacted euchromatin, allowing transcription factors to bind to regulatory sites on DNA, in order to cause transcriptional activation (22). Through the treatment with compounds function as chromatin-modifying enzyme inhibitors, we found that 5-Aza treatment enlarges the nuclear size of hESCs remarkably (Fig. 1*D*), also some other inhibitors. The great increase in nuclear size of hESCs from 5-Aza treatment indicated that modulation of chromatin-modifying enzymes is enabled to change nuclear size (Fig. 1*E*). It suggested that perturbation of chromatin-modifying enzyme activity alters nuclear size in normal cells.

**Fig. 1.**
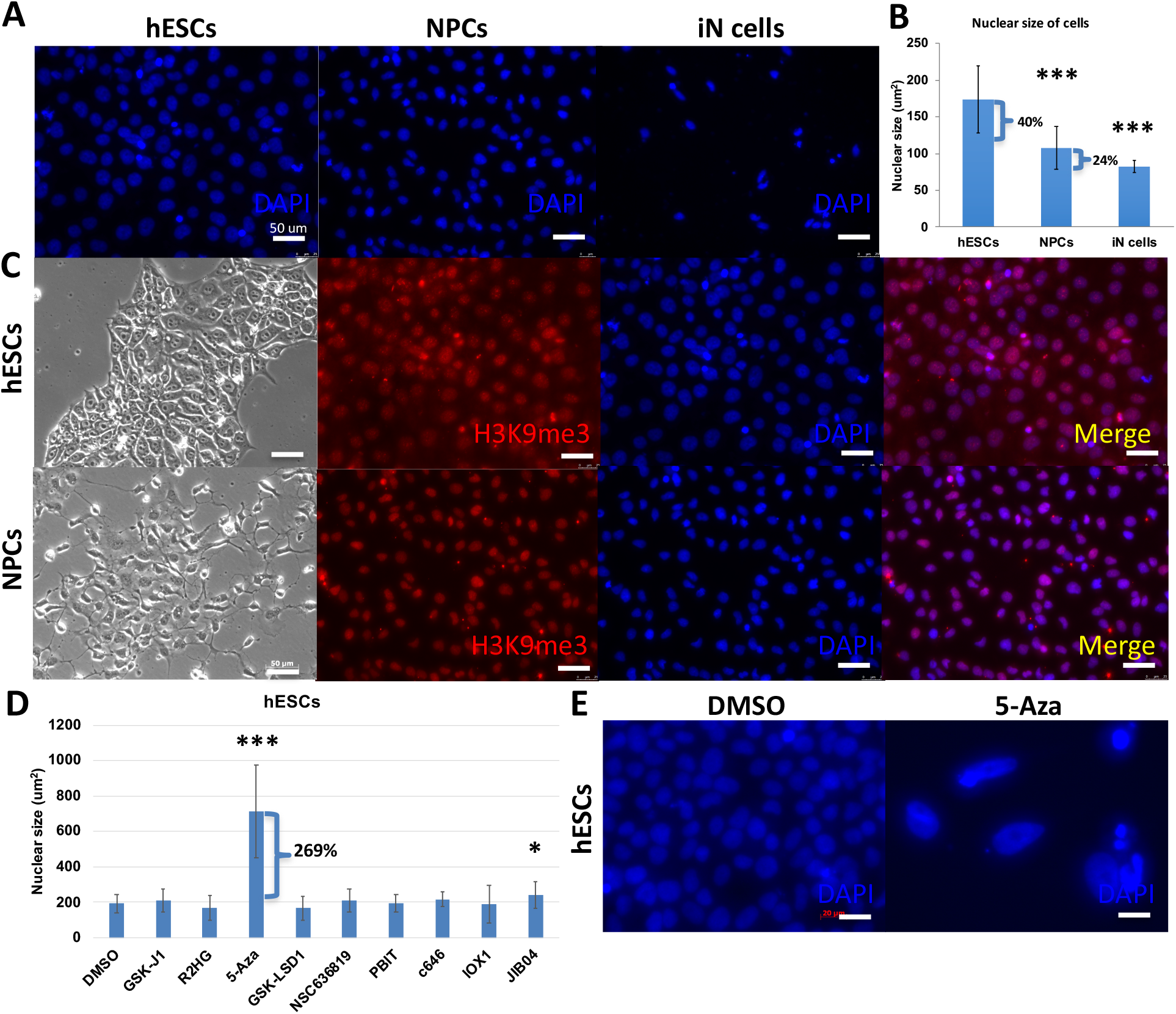
Nuclear size is associated with cell fate decision and chromatin-modifying enzyme activity. **(*A*)** Immunofluorescence staining cell nucleus of hESCs, hESC-derived NPCs, hESC-derived iN cells by DAPI (blue). White scale bars= 50 μm. **(*B*)** Measurements of cell nucleus size of hESCs, hESC-derived NPCs, hESC-derived iN cells by DAPI staining. ***P < 0.001. **(*C*)** Brightfield and fluorescent images of hESCs and hESC-derived NPCs immunofluorescence staining for H3K9me3 (red) and DAPI (blue). ***(D)*** Measurements of nuclear size after hESCs treated with histone demethylase inhibitors (GSK J1, (R)-2-Hydroxyglutaric acid disodium salt, GSK-LSD1, NSC636819, PBIT, IOX1, JIB04), DNA methyltransferase inhibitor (5-Azacytidine), and histone acetyltransferase inhibitor (C646) by DAPI staining. Statistically significant differences are marked with asterisks. *P < 0.05, ***P < 0.001. **(*E*)** Representative immunocytochemistry images of hESCs stained with DAPI. hESCs treated with DMSO as control and 5-Aza. It was shown that the bigger nuclei occurred at 5-Aza treatment. The images were taken at 40× on a confocal microscope. White scale bars= 20 μm.

In addition, we noticed that the treatment of high concentrations of IOX1 revealed the cytotoxic effects on hESCs. Therefore, we tested a series of concentrations of compounds for hESCs treatment. The high concentrations of inhibitors caused cell death of hESCs, such as 100 mM GSK-J1, 100 mM IOX1, 1 mM JIB04, 10 mM PBIT, 25 mM C646, especially, 10 μM, 1 mM, 10 mM 5-Aza induced numerous of hESCs death. The high concentrations of inhibitors have cytotoxic effects on cells and impact cells alive. However, hESCs grow well in 60 μM, 1 mM, 10 mM GSK-J1, 100 mM R2HG, 10 mM and 100 mM GSK-LSD1, 1 mM and 10 mM IOX1, 10 μM and 100 μM JIB04, 10 μM and 10 mM NSC636819, 100 μM and 1 mM PBIT, 250 μM and 2.5 mM C646 treatment. It indicated that compounds function as chromatin-modifying enzyme inhibitors which effect nuclear size alteration, possibly through modulating histone dynamics. It had been shown that utilizing histone methyltransferase inhibitors to decrease heterochromatin results in a softer nucleus and nuclear blebbing, without perturbing lamins. Histone demethylase inhibitors increase heterochromatin and chromatin nuclear rigidity (23). Given that measurements in these reports demonstrate that there is evidence that nuclear size abnormalities can be caused by alterations to chromatin regulatory proteins.

### Inhibition of chromatin-modifying enzyme effects nuclear size and cell identification of NPCs

Not only hESCs, but we also found that the treatment of 5-Aza, C646, and JIB04 enhanced the nuclear size of hESC-derived NPCs. In contrast, the treatment of R2HG, GSK-LSD1, NSC636819, IOX1 reduced the nuclear size of hESC-derived NPCs (Fig. 2*A*). 5-Aza treatment slightly reduced the nuclear size of hESC-derived iN cells (Fig. 2*B*). To gain further insights into whether these inhibitors not only impact nuclear size and cell death but also effect additional features, such as cell fate decisions, we treated the hESC-derived NPCs with DMSO and chromatin-modifying enzyme inhibitors. The concentration we used was tested and the same concentration was used as hESCs treatment, then imaged hESC-derived NPCs after staining for Sox1 (green), PI (red), and DAPI (blue) (Fig. 2*C*). As known, Sox1 is a neural stem cell marker and is highly expressed in NPCs. It was observed that the expression of Sox1 was significantly reduced in a large number of NPCs in 5-Aza, JIB04, PBIT, GSK-J1, GSK-LSD1, c646, and IOX1 treatment (Fig. 2*D*). To eliminate the possibility of Sox1 expression decrease caused by cell death, we performed propidium iodide (PI) staining to NPCs after inhibitors treatment. As known, PI is not permanent to live cells but is used to detect dead cells in a population. It was shown that Sox1 silence was not caused by cell death due to PI staining being negative. Except IOX1, JIB04 and GSK-LSD1 increased PI staining in NPCs which suggested that it may induces cytotoxic responses of NPCs occur as a consequence of cell death. Therefore, it suggested that the compound inhibitors of chromatin-modifying enzymes perturb cell identification. Collectively, inhibition of chromatin-modifying enzymes activity enabled to impact nuclear size in both hESCs and NPCs which consequently leads cell fate to aberrant states.

**Fig. 2.**
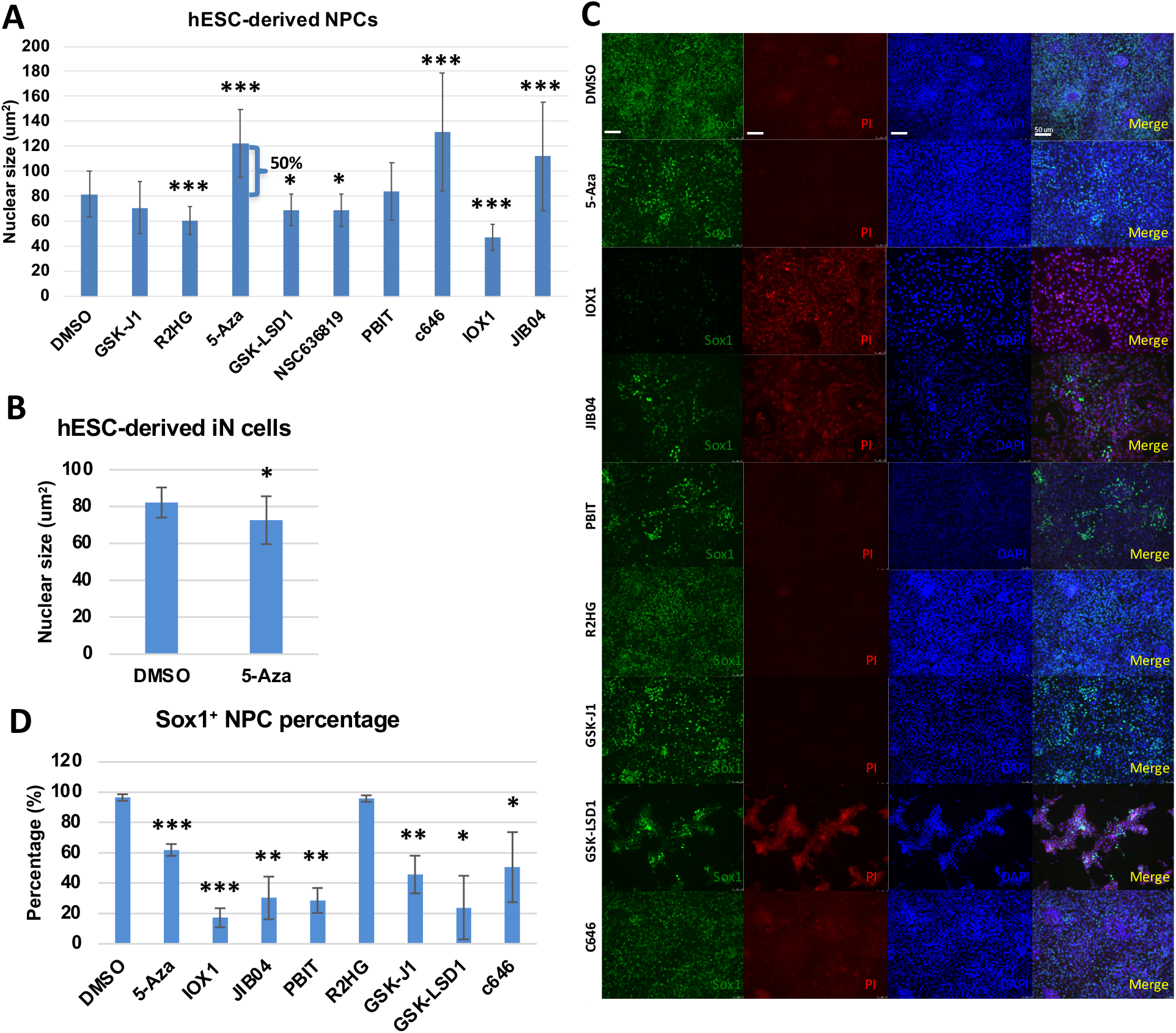
Inhibition of chromatin-modifying enzyme effects nuclear size and cell identification of NPCs. **(*A*)** Measurements of nuclear size by DAPI staining after hESC-derived NPCs treated with DMSO as control and chromatin-modifying enzymes inhibitors. **(*B*)** Measurements of nuclear size by DAPI staining after hESC-derived iN cells treated with DMSO as control and 5-Aza. **(*C*)** Fluorescent images of hESC-derived NPCs immunofluorescence staining for Sox1 (green), PI (red), and DAPI (blue) after treatment with DMSO as control and inhibitors of chromatin-modifying enzymes. White scale bars= 50 μm. **(*D*)** Percentage of cells expressing Sox1 (green) over DAPI (blue) total cells. *P < 0.05, **P < 0.01, ***P < 0.001.

### Upregulated GASC1 impairs differentiation from hESCs to iN cells

As we previously discussed, the nuclear size dramatically reduced from hESCs to NPCs, and to iN cells during lineage differentiation. Additionally, we want to validate the specificity of the targeted chromatin-modifying enzyme instead of using chemical compounds. So, we selected three represented histone enzymes such as GASC1, DNMT3A, and KDM4A, based on the screening of inhibitors of chromatin-modifying enzyme activity, to investigate the impact of chromatin-modifying enzymes in lineage differentiation. GASC1 (lysine-specific demethylase 4C, KDM4C) is a histone demethylase as well as KDM4A (lysine-specific demethylase 4A), they both belong to the families of lysine demethylases (KDM). DNMT3A (DNA methyltransferase 3A) catalyzes 5-methylcytosine methylation. 5-Aza inhibits DNA methyltransferase (DNMT). We used the doxycycline (dox)-inducible lentiviral vector system to transduce empty vector plasmid as control and GASC1, DNMT3A, KDM4A into the hESCs H1 cell line, separately. As mentioned above, we used the same approach to induce neuronal cells from differentiating hESCs by Ngn2 overexpression (20). It was shown that there is a significantly high efficiency of wt hESCs differentiating to iN cells as nearly 100%. The iN cells display a mature neural morphology with axon extension and packaged smaller cell nucleus (Fig. 3*A*). We stained hESC-iN cells with a neuronal marker Tuj1 (tubulin beta-III) and a pluripotent marker Sox2. It demonstrated that iN cells differentiated from wt hESCs showed Tuj1 positive expression and long axon extension, no Sox2 indicated hESCs left, which also occurred in KDM4A overexpressed hESC-iN cells. In contrast, hESC-iN cells with induced overexpression of GASC1 and DNMT3A that grew few extend axons and showed big nuclei. In addition, a large number of cells were Sox2 positive during differentiation (Fig. 3*A*). After measurement analysis of neurite length at 14 days of iN cell differentiation, we found that hESC-iN cells with GASC1 and DNMT3A overexpression grew remarkably shorter neurite compared to wt and KDM4A overexpressed iN cells (Fig. 3*B*). Perturbation of chromatin-modifying enzymes activity alters the molecular and cellular identity of stem cells. Overall, these data suggested that chromatin-modifying enzymes can drive lineage specification as well as assist nuclear reconstruction, in order to facilitate a smoother transition from stem cells to differentiated cells.

**Fig. 3.**
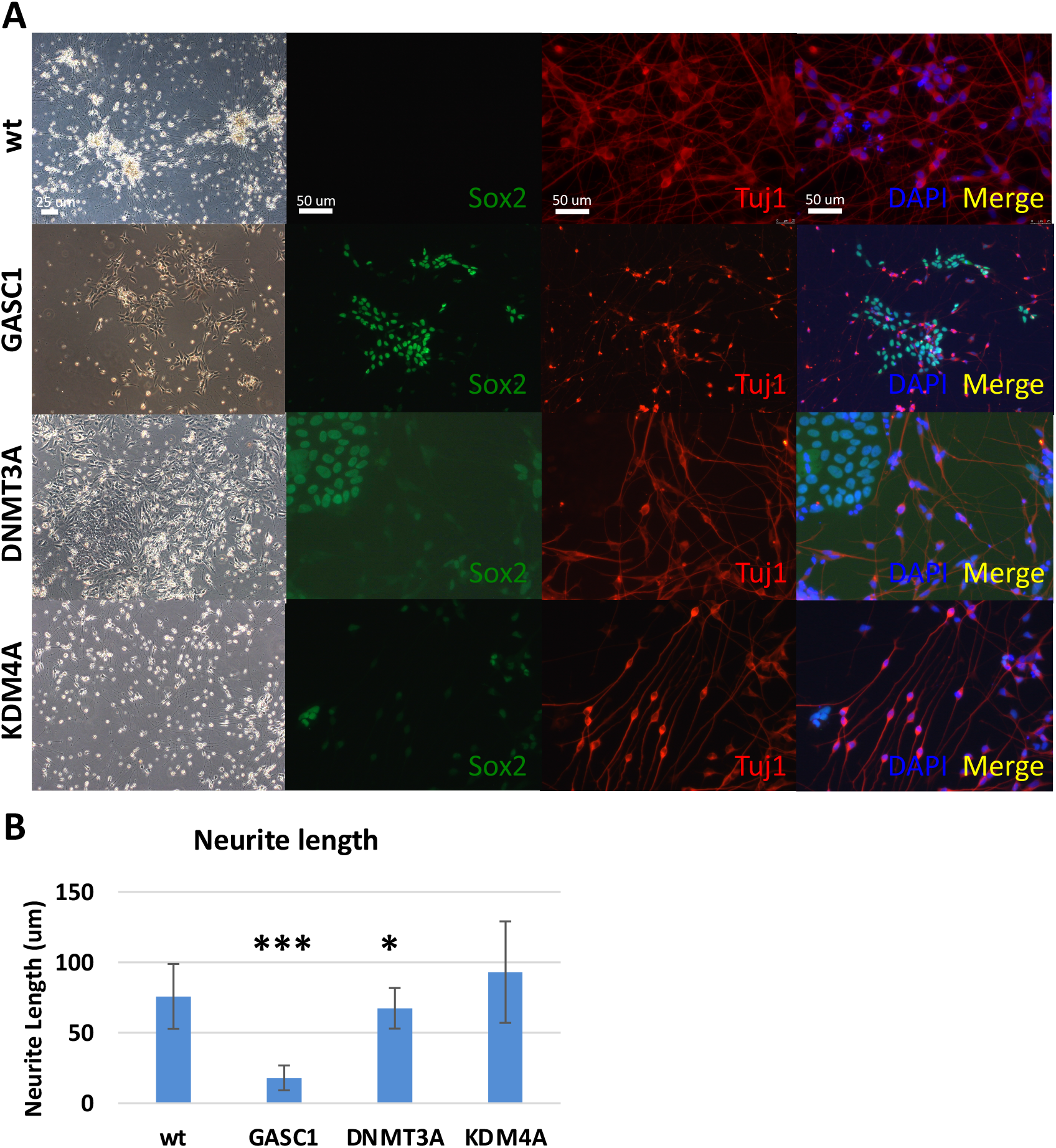
Upregulated GASC1 impairs differentiation from hESCs to iN cells. **(*A*)** Brightfield and fluorescent images of hESC-derived iN cells with wt, dox-induced overexpression of GASC1, DNMT3A, and KDM4A stained for the pluripotent marker Sox2 (green), the neural cells marker Tuj1 (red), and nucleus staining DAPI (blue) after 14 days of iN cells differentiation. **(*B*)** Neurite length of hESC-derived iN cells was measured at 14 days of iN cells differentiation. p-values were calculated by unpaired t-test, error bars represent standard deviation, *P < 0.05, *** P< 0.001.

### GASC1 promotes aberrant nuclear structure and cell cycle arrest of leukemia cells

As above discussed, chromatin-modifying enzyme activity is highly associated with nuclear size alteration in normal stem cells during lineage differentiation. As reported, cancer cells are known to develop larger nuclei as they become more malignant and increase the nuclear-to-cytoplasmic ratio (24). The mechanisms that underlie the increase in nuclear size in malignant cells remain largely unknown. Abnormal nuclear morphology has been used for cancer diagnoses for nearly a century in tests such as the Pap smear (25). We measured the nuclear size of K562 leukemia cells by DAPI staining, in order to investigate the distinctive features of chromatin-modifying enzyme activity in cancer cells compared to normal stem cells (Fig. 4*A* and *B*). We tested a series of concentrations of compound inhibitors for K562 cells treatment. Consistent with hESCs, high concentrations of inhibitors caused cell death of K562 cells. Especially, high doses and long-time of 5-Aza and PBIT treatment induce cell death. In addition, we photographed the DAPI stained nucleus of K562 cells treated with DMSO as control and 5-Aza, IOX1, GSK-LSD1, and JIB04. It was shown that K562 cells treated with inhibitors have a bigger size of nucleus and twisted shape as clover, especially in 5-Aza, IOX1, GSK-LSD1, and JIB04 treatment (Fig. 4*A*). It demonstrated that the irregularity of nuclear contour occurred in inhibitors treatment from the visible fluorescent images. The most studied types of nuclear morphological abnormalities are protrusions from the normally ellipsoidal nucleus, termed “blebs.” K562 cells showed a “blebs” shape and seem to prefer to have a twisted nuclear shape but not die as quickly as hESCs and NPCs did. It all suggested that not only nuclear size change but also nucleus misshapen are indicated the pathological change of K562 leukemia cells induced by inhibition of chromatin-modifying enzyme activity. Furthermore, the size of cancer cells was more similar to hESCs but bigger than NPCs. After inhibitors treatment for 7 days (10 μM GSK-J1, 100 μM R2HG, 1 μM 5-Aza, 10 μM GSK-LSD1, 10 μM IOX1, 100 μM JIB04, 10 μM NSC636819, 1 μM PBIT, 2.5 μM C646), compared to DMSO control, the increased percentage of nuclear size of K562 cells is 5% ∼ 44%. The treatment of 5-Aza and IOX1 markedly lead to the enlargement of the nuclear size of K562 leukemia cells (Fig. 4*B*). Except for lamina perturbation affecting nuclear abnormalities (26), it also suggested that enlarged and twisted nuclear size effect the survival of K562 cells which is associated with chromatin-modifying enzyme activity.

**Fig. 4.**
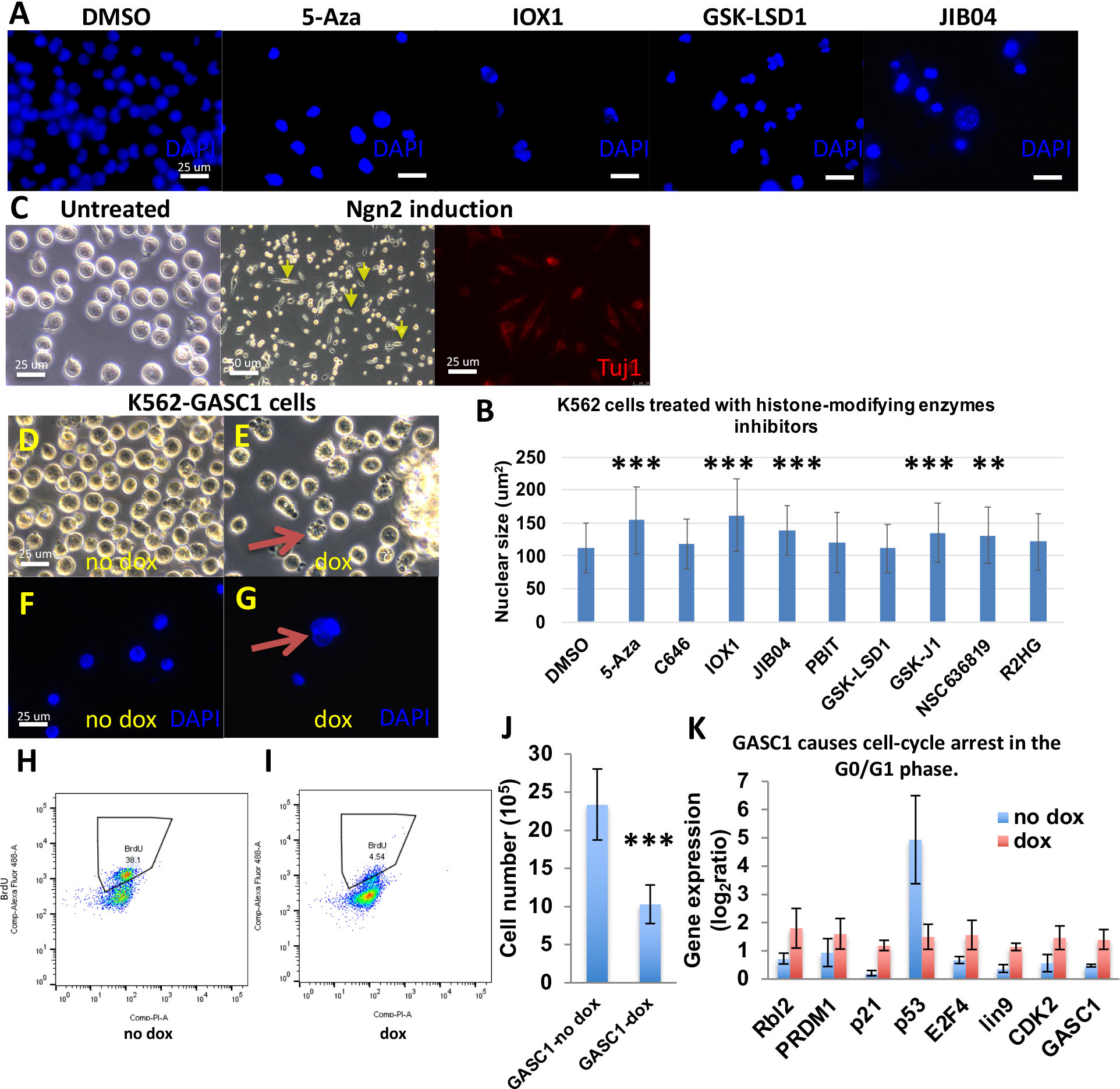
GASC1 promotes aberrant nuclear structure and cell cycle arrest of leukemia cells. **(*A*)** Representative images of DAPI staining for K562 cells treated with DMSO as control and 5-Aza, IOX1, GSK-LSD1, and JIB04. It was shown that the bigger size and twisted shape of nuclei in inhibitors treatment. The images were taken at 40× on a confocal microscope. White scale bars= 25 μm. **(*B*)** Measurements of the nuclear size using DAPI staining under fluorescence microscopy after K562 cells treated with inhibitors of chromatin-modifying enzymes. Two-tailed t-test, Mean ± SD. **P < 0.01, ***P < 0.001. **(*C*)** Brightfield and fluorescent images show that K562 leukemia cells differentiate to iN cells with Ngn2 overexpression after 14 days differentiation and stain with the neuronal marker Tuj1 (red), compared to healthy and as cultured K562 cells without any treatment. The yellow arrows indicate the cells of morphological changes. **(*D* and *E*)** Brightfield images of K562 leukemia cells with induced expression of GASC1 with or without doxycycline (dox). The no dox-induced GASC1 overexpression K562 cells show spherical morphology, same as normal cells. The dox-induced GASC1 overexpression K562 cells show morphology change from spherical to twisted shape and cell blebs. The red arrow indicates the cells with twisted shapes and blebs. White scale bars= 25 μm. **(*F* and *G*)** Fluorescent images by DAPI staining demonstrate that the dox-induced GASC1 overexpression K562 cells show the twisted nuclear shape and bigger nuclei compared with the no dox-induced GASC1 overexpression K562 cells. The red arrow indicates the enlarged and twisted nucleus. White scale bars= 25 μm. **(*H* and *I*)** Flow cytometric analysis of K562 cells were double-stained with BrdU and PI. The dox-induced GASC1 overexpression K562 cells proliferate significantly slower than the no dox-induced GASC1 overexpression K562 cells by BrdU assay. **(*J*)** Cell number counting after induction by dox for 7 days. The cell number is decreased in dox-induced GASC1 overexpression K562 cells than no dox-induced GASC1 overexpression K562 cells. ***P < 0.001. n=3. **(*K*)** The genes play a role in cell-cycle arrest detected by qPCR in the dox-induced GASC1 overexpression K562 cells and no dox-induced GASC1 overexpression K562 cells. Unpaired two-tailed Student t-test. Data are mean ± SD. P < 0.001.

As mentioned above, these chromatin-modifying enzymes effect lineage differentiation of hESCs. These hESC-iN cells with GASC1 and DNMT3A aberrant expression retain their proliferation capability, fail to differentiate, and acquire resistance to cell death (Fig. 3). As previously reported, IDH1 mutated cells generate R(2)-2-hydroxyglutarate (2HG) (27), which causes GASC1 mediated hypermethylation (15, 28). IDH1 gene mutations occur frequently in AML (acute lymphoblastic leukemia). Accordingly, we investigated the role of GASC1 in K562 leukemia cells differentiation. Basically, healthy K562 cells are spherical in shape without any structural distortions, and grow in suspension (Fig. 4*C*). After being differentiated by Ngn2 overexpression for 14 days, it was shown that a portion of K562 cells grew spike, and Tuj1 was not abundantly expressed (Fig. 4*C*). K562 cells are hard to differentiate. When adding dox in cell culture for inducing GASC1 overexpression, the majority of K562 leukemia cells began to change their morphology from smooth surfaces of spherical shape to twisted cellular shape compared with no dox-induced K562 cells (Fig. 4*D* and *E*). The abnormal nuclear morphology and protrusions termed “blebs” are both associated with chromatin alterations (23). To validate the cell shape change corresponding to nucleus change, we stain K562 cells with DAPI. Under microscope examination of GASC1 overexpressed K562 cells with or without dox, we found that it is consistent with cell shape change in dox-induced overexpressed K562 cells, that showed nucleus enlarges and shapes as clover (Fig. 4*F* and *G*). To investigate whether the twisted shape consequently provides benefits of growth of the K562 cells or not, we employed the BrdU cell proliferation assay to interrogate the cell cycle of K562 cells growth. The dox-induced K562-GASC1 cells significantly proliferate slower than the no dox-induced K562-GASC1 cells (Fig. 4*H* and *I*). Meanwhile, we count the cell number of K562 cells with or without dox-induced GASC1 overexpression, it reveals that there is a significant reduction in the growth of K562 cells with GASC1 overexpression (Fig. 4*J*). To explore the mechanism of K562 cells with GASC1 expression was able to stop proliferation at the genetic level, we selected a range of key regular genes function as cell cycle effectors, such as Rbl2, PRDM1, p21, p53, E2F4, lin9, and CDK2 (29, 30, 31). Based on the qPCR results, we found that there is a remarkably increased transcription level of Rbl2, PRDM1, p21, E2F4, lin9, and CDK2 in the dox-induced GASC1 expression K562 cells than in no dox cells (Fig. 4*K*). p53 mRNA level was reduced markedly in the dox-induced GASC1 expression K562 cells. As known, depletion of p53 in cells causes G1 arrest and apoptosis. Taken together, modulation of GASC1 expression in leukemia cells leads to cell cycle arrest in the G0/G1 phase and shows misshapen nuclei. Through regulating the activity of chromatin-modifying enzymes, that lead to nuclear rupture and blebs, we sought to address a novel strategy to regulate cancer cell growth.

## Discussion

The objective of this study was to investigate the manner in which both cellular and nuclear size change during differentiation and how such changes might relevant to chromatin-modifying enzymes. However, to date, there have not been many investigations of chromatin contribution to nuclear morphology, although altered histone modifications and chromatin decompaction are consistently present in both nuclei with abnormal morphology and nuclear blebs.

Lamins are fibrous proteins as well as nuclear structural proteins. Mouse ES cells express high levels of A-type lamin during cell differentiation (32, 33). Forced expression of the progerin mutant isoform of lamin A/C (LMAC) increases osteogenic differentiation and reduces adipogenic differentiation (33). Additionally, deregulation of lamin expression can lead to various cancer and be used as diagnostic biomarkers. Lamin is a genetic reason for nuclear misshapen, however epigenetic mechanism of nuclear shape change in cell fate determination is still obscure. Here, we aim to understand whether activity alterations of chromatin-modifying enzymes affect nuclear contour, and what mechanisms contribute to cellular identification of cell fate. We employed histone-modifying enzyme inhibitors to get some clues of whether certain types of enzymes or all histone-modifying enzymes are able to effect nuclear size alteration. We used stem cells such as hESCs and NPCs to investigate the impact of histone-modifying enzyme inhibitors at the stage of pluripotency and multipotency maintenance, which may have some distinctions. Also, we investigated the impact of histone-modifying enzyme inhibitors on cancer cells such as leukemia cells. We observed that there is enlarged nuclear size and misshapen nucleus via chromatin-modifying enzyme inhibition. As known, heterochromatin appears larger, fewer, and decondensed in ES cells, numerous and condensed in ES cell-derived neuronal progenitor cells, and appear fewer but more compact in terminally differentiated neurons (34, 35). Additionally, cancer cells have global alterations in chromatin organization, mostly a reduction in heterochromatin (36). Therefore, it suggested that the original histone states identify the specific type of cell, and impact the outcome of histone-modifying enzymes inhibitors treatment.

In addition, when treated with histone-modifying enzyme inhibitors, the nuclear size was able to alter, and the nuclear shape becomes aberrant. It demonstrated that chromatin state is a major factor determining the morphology of the nucleus. The increase of nuclear blebbing in nucleus which indicated that a consequence of increasing euchromatin or decreasing heterochromatin, and decreasing nuclear rigidity. In addition, during dysregulation of histone by treatment of histone-modifying enzyme inhibitors, some cells were unable to endure the stimulation and die, but some cells with a misshapen nucleus survived. Is that the reconstructive chromatin provides certain benefits for cell survival during chromatin-modifying enzyme dysregulation? After treatment of histone-modifying enzyme inhibitors in NPCs, it was shown that the decreased expression of genes who play a role in stem cell identification. We also observed the nuclear size become smaller and smaller when cells were differentiated from hESCs to NPCs, then to iN cells. Simultaneously, chromatin appears decondensed in ES cells, then appears more compact in differentiated neurons. Collectively, it implies that the major function of chromatin-modifying enzyme is to determine cell fate, nuclear size alteration is a phenomenon accompanied by the occurrence of cell lineage commitment in normal cells.

As known, histone modifications correlate with ESC differentiation. Hundreds of promoters of pluripotent genes gain their epigenetic marks, whereas many others lose it, during the transition from ESC to NPCs, and from NPCs to differentiated neurons (37, 38). Through modulating chromatin-modifying enzymes that are able to control a huge number of epigenetic marks on the promoters of stem cell genes. Therefore, we ask for is that possible to modulate chromatin-modifying enzymes to alter cancer cell fate. We selected three representative chromatin-modifying enzymes. Unlike KDM4A-B, KDM4C (GASC1) is associated with chromatin during mitosis, and triggers the demethylation of H3K9me3 mark (10). Ectopic expression of GASC1 or other JMJD2 members markedly decreases H3K9me3/me2 levels, increases H3K9me1 levels, delocalizes HP1 (heterochromatin protein 1), and reduces heterochromatin in vivo (15). GASC1 is an Oct4 target in mouse ES cells, and acts as a positive regulator of Nanog, also is preferentially expressed in undifferentiated ES cells (8, 39). Depletion of GASC1 in ES cells induced differentiation with a global increase in H3K9me3, confirming that the histone demethylase activity of GASC1 is linked to the maintenance of pluripotency. The expression levels of GASC1 were also significantly related to the grade of basal-like breast cancers which is the most aggressive type (14). It suggested that histone-modifying enzymes impact the cell fate of normal cells as well as cancer cells. To investigate the impact of GASC1 in cancer cells, we manipulated GASC1 expression in a commonly used leukemia cell line K562. We found that overexpression GASC1 impairs the cell cycle process, deprives K562 cells proliferation capacity, and shows nuclear abnormalities. It indicated that chromatin-modifying enzymes are associated with nuclear size determination and cell fate decision of normal cells as well as cancer cells. Through modulating chromatin-modifying enzymes activity which enables to deprive the aggressive proliferation capacity of cancer cells, arrest cells in the cell cycle to die, and as a consequence, show nucleus misshapen.

## Methods

### Cell culture and media

All cell types were maintained at 37°C in a humidified 5% (vol/vol) CO2 environment during culture. H1 ES cells were obtained from WiCell Research Resources (WiCell, WI). ES cells were maintained as feeder-free cells in mTeSR™1 medium (Stem Cell Technologies). Cells were allowed to spread overnight prior to transduction. We used a doxycycline (dox)-inducible lentiviral vector system to transduce empty vector plasmid as a control and GASC1, DNMT3A, KDM4A into the H1 ES cells, separately. Selection was carried out for 7 days in mTeSR™1 medium with 50 μg/ml hygromycin.

### Generation of iN cells from human ES cells

H1 human ESCs were dissociated using accutase and plated as single cells in mTeSR™1 medium supplemented with ROCK inhibitor Y-27632 (10 μM, StemCell Technologies, Inc.) on matrigel to a final density of 20,000-30,000 cells per cm^2^. On the second day, attached cells were infected with lentivirus for Ngn2-puro and rtTA expression. After 24 hr (day 0), transgene expression was induced by replacing the medium with N2/B27 medium (DMEM/F12, N2 supplement (Life Technologies), B27 supplement (Life Technologies), 10 μg/ml insulin (Life Technologies)), and 2 μg/ml Doxycycline. Selection was carried out for 14 days (day 1 to day 14) in N2/B27 medium with 2 μg/ml puromycin. From day 2, induced neuronal cells were maintained in N2/B27 medium and changed medium every 2 days.

### hESCs differentiate to neural progenitor cells

Growing hESCs in mTeSR media on matrigel coated dish until confluent, then enzymatically passage with accutase, resuspend in Neurobasal-A medium, DMEM/F12, 1% N2, 1% B27 and insulin media, with Y-27632, a specific inhibitor of Rho kinase activity, cultured in the same size of the dish without matrigel coated. Overnight, colonies will form floating spherical clusters termed “embryoid bodies” (EBs). Wait until the EB size grows to 200∼400 μm, gently move it to matrigel coated dish. The next day, it will attach to the bottom. Changing medium with N2/B27 media supplemented with dual SMAD inhibitors (LDN193189 and SB431542). Feed EBs every second day with N2/B27 media supplemented with 0.1 μM LDN193189 and 10 μM SB431542. After several days, the neural rosettes begin to appear, characterized as round clusters of neuroepithelial cells with apicobasal polarity. Staining Sox1 to indicate the neural progenitor cells.

### Virus generation

Lentiviruses were produced as described (40) in HEK293T cells (ATCC, VA) by co-transfection with three helper plasmids (pRSV-REV, pMDLg/pRRE and vesicular stomatitis virus G protein expression vector) (12 μg of lentiviral vector DNA and 6 μg of each of the helper plasmid DNA per 75 cm^2^ culture area) using calcium phosphate (41). Lentiviruses were harvested with the medium 24 and 46 hr after transfection. Viral supernatant was filtered through a 0.45 μm filter, pelleted by centrifugation (21,000 × g for 2 hours), resuspended in DMEM, aliquoted, and snap-frozen in liquid N_2_. Only virus preparations with >90% infection efficiency as assessed by EGFP expression or puromycin resistance were used for experiments.

### Lentiviral infections

Complementary DNAs for candidate genes were cloned into doxycycline-inducible lentiviral vectors, as described previously (42). Lentiviral production and cell infection were performed as described previously (43). Cells were infected with concentrated lentivirus and treated with doxycycline (2 mg/ul) 16-24 h later.

### Immunofluorescence

Cultured iN cells were fixed in 4% paraformaldehyde in PBS for 15 min at room temperature, washed three times with PBS, and incubated in 0.2% Triton X-100 in PBS for 10 min at room temperature. Cells were blocked in PBS containing 10% BSA for 3 hours at 4°C. Primary antibodies were applied overnight. Cells were washed in PBS three times. Secondary antibodies were applied for 3 hours, washed. 300 nM DAPI staining for indication of nucleus. Mount.

### Antibody

The following primary antibodies with indicated dilution in blocking buffer were used: Rabbit anti-Tuj1 (Covance, BioLegend, MRB-435P, 1:1,000), Mouse anti-Tuj1 (Covance, BioLegend, MMS-435P, 1:1,000), Rabbit anti-human Sox1 (Millipore, AB15766, 1:200), Goat anti-Sox2 (Santa Cruz, 1:200). After staining with corresponding secondary antibodies in PBS plus 1% BSA, coverslips were washed four times with PBS, each for 10 min, mounted with the mounting medium onto glass slides, and examined under Olympus FluoView FV1000.

### Inhibitors of chromatin-modifying enzymes

Purchase from tocris. GSK J1 (Cat. No. 4593), (R)-2-Hydroxyglutaric acid disodium salt (R2HG, Cat. No. 6124), GSK-LSD 1 dihydrochloride (Cat. No. 5361), IOX 1 (Cat. No. 4464/10), JIB04 (Cat. No. 4972/10), NSC636819 (Cat. No. 5287/10), PBIT (Cat. No. 5930), C646 (Cat. No. 4200/10), 5-azacytidine (5-Aza, Cat. No. 3842).

### qRT-PCR

RNA was isolated through standard methods following manufacturer’s instructions (QIAGEN), and reverse transcribed with Superscript III (Life Technology). qRT-PCR was performed with Fast Start universal SYBR green master mix (Roche) along with gene specific primers on a 7900HT real-time PCR machine (Applied Biosystems). Statistical significance was determined by Students t test based on triplicated experiments.

### Statistical Tests

Microsoft Excel was used for data analysis. Cumulative plots were generated from parameters collected from individual cells under similar experimental conditions. Average data are presented as bar graphs indicating means ± SD. Statistical comparisons between bar graphs were made using the unpaired, two-tail, Student t test (*P < 0.05, **P < 0.01, and ***P < 0.001, all versus control).

## Acknowledgements

I would like to thank all our laboratory members and collaborators for their comments and dedicated work that have greatly contributed to the ideas presented here.

## Author Contributions

L.M. conceived and designed the study; L.M. planned and performed the experiments; L.M. wrote the manuscript.

